# Non-academic employability of life science PhDs: the importance of training beyond the bench

**DOI:** 10.1101/485268

**Authors:** Sohyoung Her, Mathieu D Jacob, Sharon Wang, Songyi Xu, David CF Sealey

**Affiliations:** Department of Pharmaceutical Sciences, University of Toronto, Ontario, Canada; Life Sciences Career Development Society, University of Toronto, Ontario, Canada; Science Career Impact Project, Ontario, Canada; Department of Laboratory Medicine and Pathobiology, University of Toronto; Toronto, Ontario, Canada

## Abstract

To better understand how PhD graduates have prepared for the non-academic job market, we surveyed life science PhD and postdoctoral graduates from the University of Toronto who were employed in non-academic sectors. We also surveyed life science PhD and postdoctoral trainees to assess their engagement in career preparation activities. PhD professionals employed in non-academic sectors had engaged in various career preparation activities during their training. Some activities had a higher perceived impact on the path to employment than others. Trainees had also engaged in such activities, but those rated by professionals as having a highly positive impact on their path to employment were engaged in by only a minority of trainees. The proportion of trainees who wished to work in a non-academic sector was higher among those who were closer to program completion. Like professionals, many trainees reported facing barriers to pursuing career development activities. Our findings suggest that PhD trainees seeking to work in non-academic sectors should engage in career preparation activities, particularly those that involve experiential learning. By supporting co-curricular programming and reducing barriers to participation in career development activities, academic administrators and faculty have the opportunity to support trainees’ professional development beyond the laboratory.

## INTRODUCTION

In today’s knowledge economy, PhD degrees are being awarded at an increasing rate, and the career outcomes of PhD graduates are of critical interest. While the traditional career track of PhD graduates has long been considered a research-intensive career in academia, PhD graduates are increasingly finding their way into non-academic careers.

According to a US National Science Foundation report on the labor force of science and engineering graduates from 1993 to 2010, the majority of PhD recipients were employed in sectors outside of academia including business and government (National Science Foundation 2014). In fact, over the years examined, the percentage of biological, agricultural and environmental life sciences doctorate recipients holding tenure or tenure-track appointments at academic institutions decreased from 9.0% to 7.6% for those within 3 years of graduation, and from 17.3% to 10.6% for individuals after 3 to 5 years of obtaining their degree (National Science Foundation 2014). Similarly, a study from the Higher Education Quality Council of Ontario found that as of 2015 only 10% to 15% of science PhDs who graduated in 2009 held academic professor positions, and at least 40% were employed in non-academic sectors (Jonker 2016). This is further reflected in a recent report from the University of Toronto; in 2016 over 40% of life science PhD graduates from 2000 to 2015 were employed in a non-academic sector (Reithmeier et al. 2018). These results are comparable to those at the University of British Columbia for PhD graduates from 2005 to 2013 (Porter et al. 2017). These trends reflect the imbalance between the number of PhD graduates and the availability of academic faculty positions, as well as the interest of contemporary PhD graduates in non-academic careers (Fuhrmann et al. 2011; Ghaffarzadegan et al. 2015; Larson et al. 2014; National Science Foundation 2015; Sauermann and Roach 2012; Sekuler 2014).

Despite the body of evidence documenting the employment of PhDs in non-academic sectors, the paths to non-academic professions are not well known within academia, and there is little consensus on the role of university departments in preparing trainees for such careers (Tilghman et al. 2012). As a response to the need for mentorship and training, some institutions have undertaken efforts to modernize the PhD program (e.g., www.nihbest.org; Lee and Reithmeier 2013), and student and alumni-led career training initiatives have emerged (e.g., www.gmcacanada.com; www.lscds.org; sciencecareerimpact.org; Freeman, 2017). However, cohort-level data on how PhD trainees have transitioned from academia to non-academic careers are lacking. Furthermore, data on the most impactful training activities in facilitating the transition from academia to non-academic employment are needed for the benefit of trainees, faculty, and academic administrators.

We surveyed life science PhD and postdoctoral graduates from the University of Toronto and affiliated research institutes who are employed in a non-academic sector. We also surveyed doctoral and postdoctoral trainees to assess their level of engagement in career preparation activities. By investigating the experiences and perspectives of trainees and alumni professionals, we aim to educate trainees on how alumni have prepared themselves for non-academic careers, and provide faculty and academic administrators with actionable insights on career training and development opportunities to meet the needs of today’s graduate trainees.

## METHODS

### Data collection

We developed a survey (Supplementary Information 1) and deployed it on the internet using SurveyGizmo. Target survey participants included: 1) life science PhD students and postdoctoral fellows at the University of Toronto and its affiliated research institutes (trainees), and 2) individuals who completed life science PhD programs and/or postdoctoral fellowships at the University of Toronto and affiliated research institutes after January 1st, 2010 and are employed in a non-academic sector (professionals). Trainees were invited to participate in the survey using e-mail lists from the Graduate Life Sciences Education office, affiliated research institutes and the Life Sciences Career Development Society (LSCDS) at the Faculty of Medicine, University of Toronto. Professionals were invited to participate using the e-mail list from the Advancement Office at the Faculty of Medicine, University of Toronto, and the alumni list of LSCDS. The survey was promoted on social media through LSCDS Twitter and Facebook accounts, and the Science Career Impact Project Twitter account. The authors also solicited participation among their academic and professional networks. The survey was launched on January 30, 2017 and closed to responses on March 3, 2017.

Participation in the survey was voluntary. Survey participation was incentivized by offering a chance to receive one of three gift cards with a value of $100 CAD. To participate in the draw, participants were required to provide an e-mail address; however, e-mail addresses provided for the draw were not linked to survey responses during data analysis. Participants were asked to provide an e-mail address if they were willing to answer additional questions from the authors about their responses.

### Data processing/analysis/validation

A total of 563 complete responses were received and exported from SurveyGizmo into Microsoft Excel for analysis. The following types of responses were removed from the dataset: duplicate responses (28, 5.0%) identified by e-mail addresses that were provided voluntarily; responses from professionals who self-identified as being employed in academia (43, 7.6%) which did not meet the inclusion criteria; responses from participants who otherwise did not meet the inclusion criteria. Responses from participants holding professional degrees (8, 1.4%; e.g., MD, DDS, MBA) were analyzed separately. A total of 446 complete responses were included in the final analysis.

## RESULTS

### Survey participants

We surveyed life science PhD students and postdoctoral fellows at the University of Toronto and its affiliated research institutes, as well as life science PhD and postdoctoral alumni working in non-academic sectors who completed their programs in 2010 and beyond. The total number of responses included in the final dataset was 446 (see Methods), including responses from 244 PhD candidates, 86 postdoctoral fellows, 79 alumni who completed a PhD as their highest level of training, and 37 alumni who completed a postdoctoral fellowship (Fig. 1). We estimate that the survey reached approximately 3000 PhD students, for a corresponding response rate of approximately 8%. The number of PhD and postdoctoral fellow alumni in the target group could not be determined; we expect that the response rate was less than 10%.

**Fig. 1.**
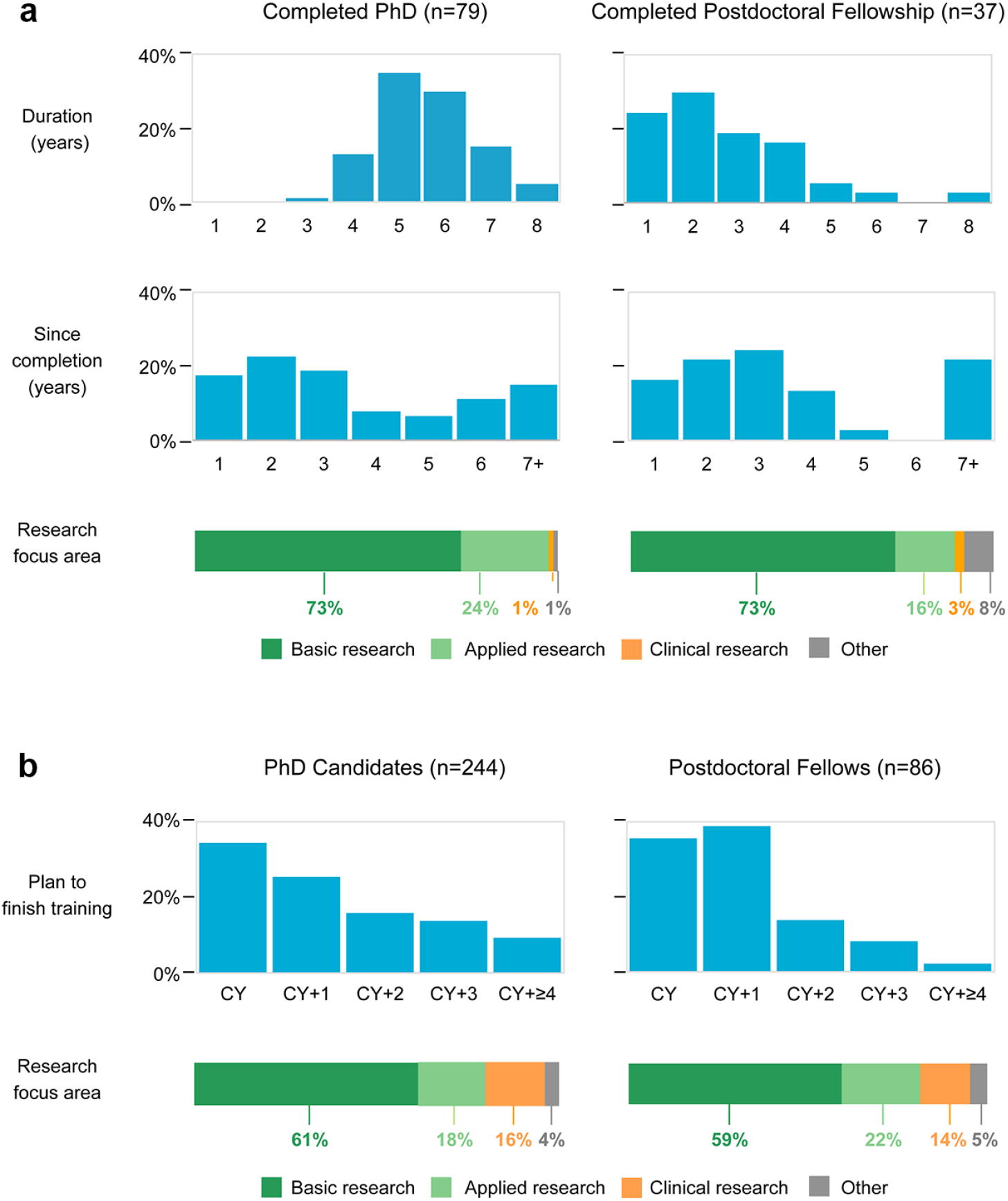
Characteristics of survey participants. **(a) Professionals.** Participants were asked about the duration of their PhD or postdoctoral fellowship (depending on their highest level of training), as well as how long ago that training was completed. Participants were asked about the focus area of their research. **(b) Trainees.** Participants were asked when they plan to finish their training, as well as the focus area of their research. CY=current year=2017.

Trainees included PhD students and postdoctoral fellows from all stages of training across multiple life science graduate departments at the University of Toronto (Fig. 1b, Supplementary Information 2). Professionals included both recent graduates and more established professionals (Fig. 1a). The mean time since completion of training was 3.5 years for professionals. Professionals were employed in a wide variety of non-academic fields; the most common areas (>10%) were research and development, medical affairs, regulatory affairs, business development, and medical/scientific communications (Fig. 2a). The mean duration of training was 5.6 years for PhD graduates and 2.7 years for completed postdoctoral fellows (Fig. 1a). The majority of survey respondents (73% of professionals; 60% of trainees) were engaged in basic research during their training (Fig. 1).

**Fig. 2.**
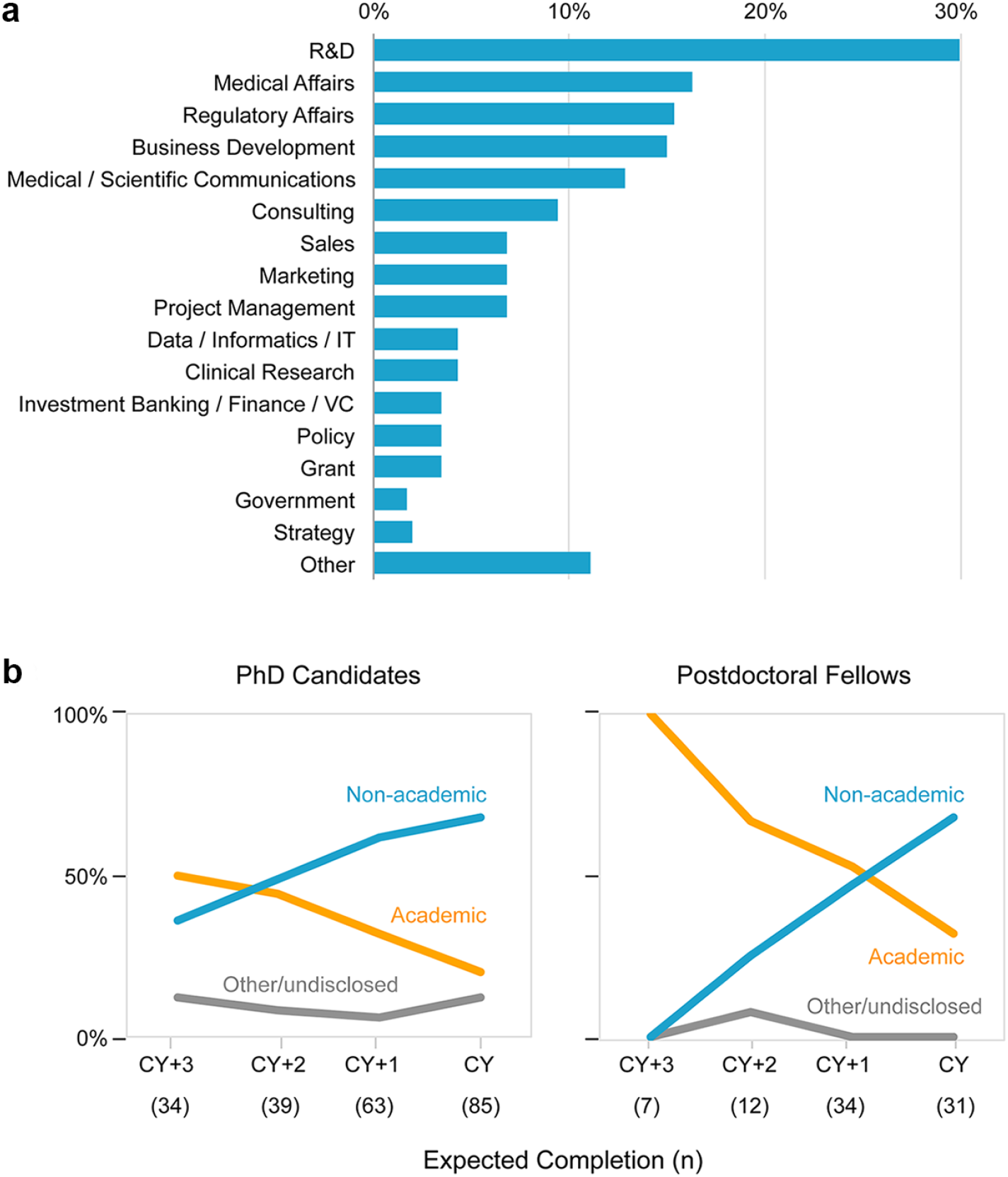
Areas of employment. **(a) Professionals.** Participants were asked what functional areas or departments they have worked in. ‘Other’ includes: corporate leadership, innovation, engineering, scientific evaluation, company incubation, operations, research administration, stakeholder relations, quality assurance, law, technology transfer and commercialization, and healthcare administration (1% each). **(b) Trainees.** Participants were asked what sector they wish to work in. The following options were provided: non-academic (e.g., industry, government, non-profit), academic (e.g., university, research institute), and other. CY=current year=2017.

Fifty-seven percent of PhD trainees wished to work in a non-academic sector (e.g., industry, government, non-profit) compared with 34% who wished to work in an academic sector (e.g., university, research institute). Forty-seven percent of postdoctoral fellows wished to work in a non-academic sector compared with 52% who wished to work in an academic sector. Interestingly, the proportion of trainees who wished to work in a non-academic sector was higher among those who were closer to program completion than those who were earlier in their programs (Fig. 2b). Sixty-eight percent of PhD candidates and postdoctoral fellows who planned to finish their training in the current year (2017) wished to work in a non-academic sector.

### Participation in and perceived importance of career development training and activities

To investigate the career development activities undertaken by professionals and trainees, as well as the perceived importance of these activities on the path to employment, we asked participants to select from a list the types of training and activities they participated in, and then to rate the impact those activities had (for professionals) or will have (for trainees) on their path to employment. We developed this list of training and activity types based on our awareness of their existence within the University of Toronto community and in professional settings.

Most respondents (97% of professionals; 96% of trainees) had participated in at least one type of career development training or activity (Fig. 3). Among professionals, the activities with the highest levels of participation (more than 50%) were career seminars, networking events, resume training, informational interviews, engagement in not-for-profit organizations or student groups, and independent study or reading. Of these, resume training, engagement in not-for-profit organizations or student groups, informational interviews, and independent study or reading were rated by more than 25% of professionals as “highly positive - no job without it.” Among the activities rated as most impactful, the top ones were internship and certificate/accreditation; however, a minority of professionals participated in these activities. Other activities rated by more than 25% of professionals as high-impact include interview training, consulting, mentorship/career coaching, and entrepreneurship. Overall, the rate of participation in the highest-impact activities varied. Almost all types of training and activities were rated by at least one professional as having a highly positive impact on the path to employment. Almost all types of training and activities were rated by more than 50% of professionals as having a highly positive or positive impact.

**Fig. 3.**
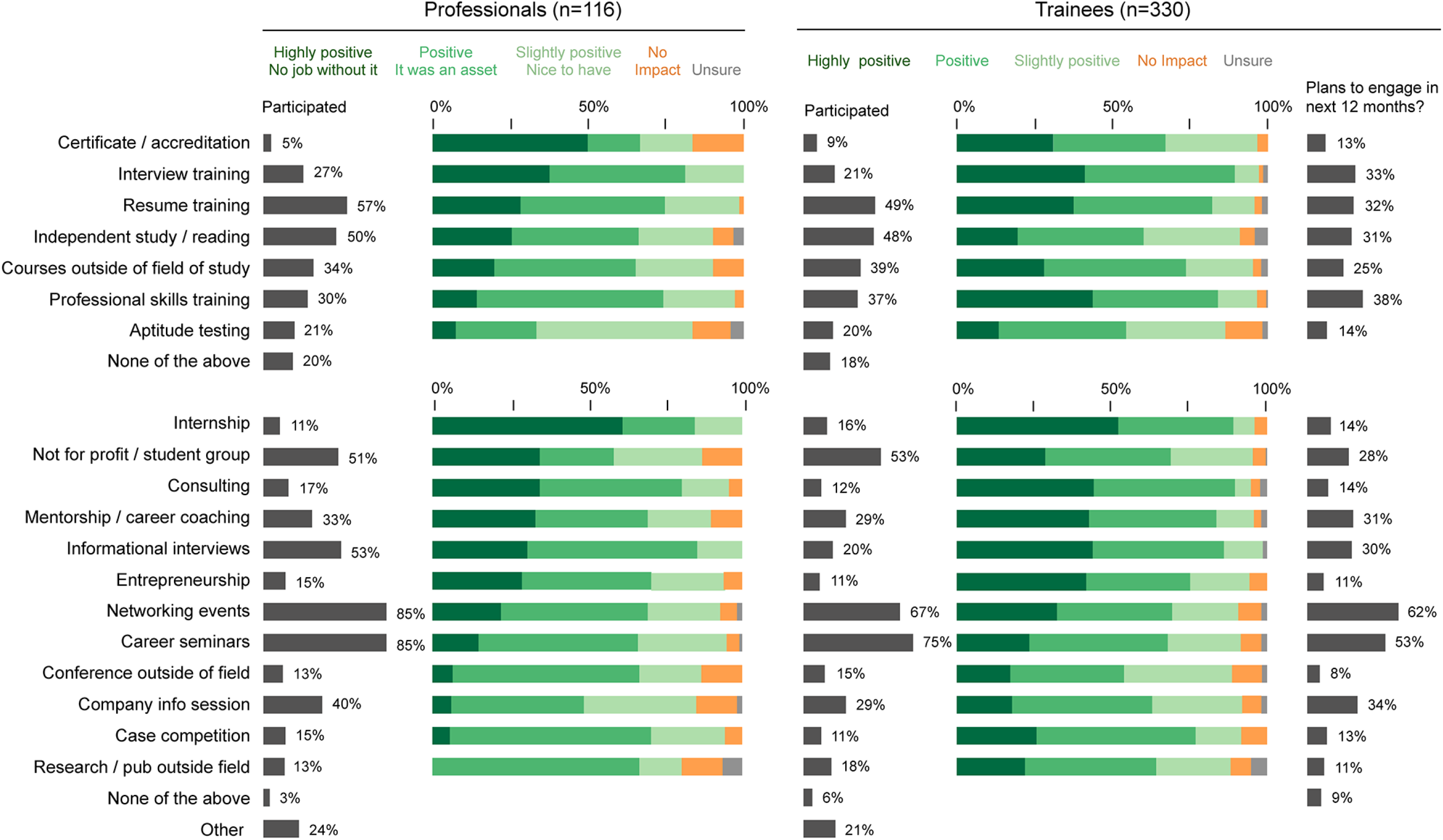
Perceived impact of extracurricular training and activities on the path to employment. **Professionals:** Participants were asked if during academic training they participated in the listed types of training (upper panel) and activities (lower panel). Participants were then asked to rate the impact their selected activities had on their path to employment. **Trainees:** Participants were asked if they participated in the listed types of training (upper panel) and activities (lower panel). Participants were then asked to rate the impact they thought these activities will have on their path to employment. Participants were asked if they planned to engage in the listed extracurricular training and activities in the next 12 months.

Overall, the rate of participation in training and activities was similar among professionals and trainees (Fig. 3). The activity with the highest disparity in participation rate was informational interviews (53% of professionals; 20% of trainees). Fifty percent of professionals engaged in at least one activity rated as “highly positive - no job without it” and 57% of trainees engaged in at least one activity that they thought would have a “highly positive” impact on their path to employment (data not shown). The perceived importance of various training and activities as rated by trainees was mostly comparable to that of professionals (Fig. 3). One exception was professional skills training, which was rated as highly positive by 44% of trainees but only 14% of professionals (Fig. 3).

Most trainees (92% of PhD candidates; 82% of postdoctoral fellows) planned to engage in extracurricular training and activities within the next 12 months; the top choices (more than 50%) were networking events and career seminars (Fig. 3). Notably, these activities were rated by only 22% and 15% of professionals, respectively, as having a highly positive impact on their path to employment.

To identify potential opportunities to develop career training programs, we compared professionals’ highest impact ratings of training and activities with the corresponding trainee participation rate (Fig. 4). Training and activities rated by more than 25% of professionals as having a highly positive impact (“no job without it”) but with trainee participation rates below 50% include certificate/accreditation, interview training, resume training, independent study/reading, internship, engagement in not-for-profit organization or student group, consulting, mentorship/career coaching, informational interviews, and entrepreneurship (Fig. 4).

**Fig. 4.**
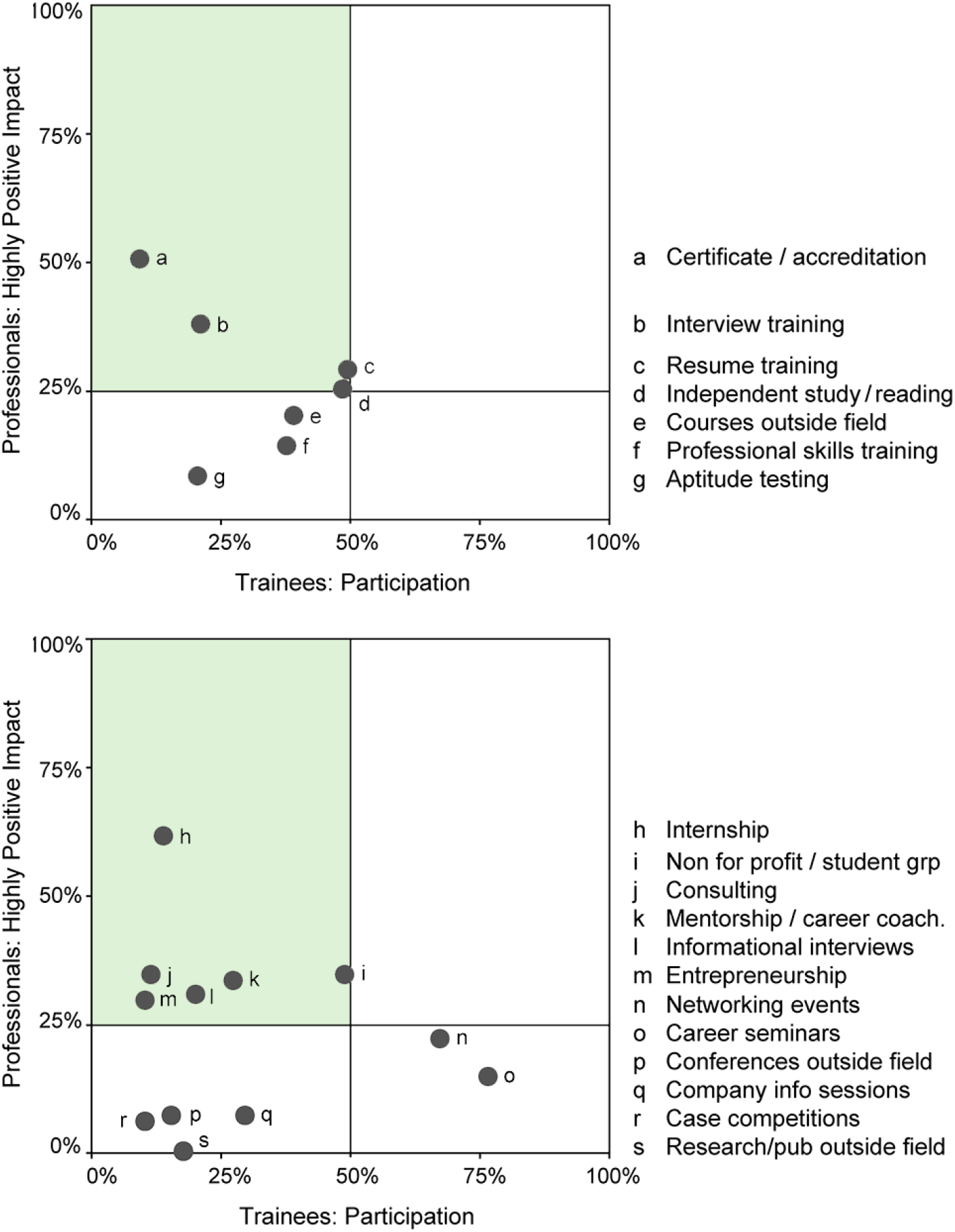
Development areas of opportunity for trainees. To visualize opportunities for trainee development, the professionals’ highest impact ratings of training and activities were plotted versus trainees’ participation rate (see Fig. 3 for data). Horizontal gridlines represent the median rating. Vertical gridlines represent 50%.

Participants reported other types of training and activities beyond the survey options including teaching/tutoring (6% of professionals; 4% of trainees), media/communications (5% of professionals; 1% of trainees), public speaking (2% of professionals; 0.6% of trainees), and athletics/art/performance (0.9% of professionals; 5% of trainees).

### Barriers to career development

Anticipating that the rate of participation in career development training and activities would vary, we asked participants if they faced (during their academic training for professionals) or currently face (for trainees) any barriers to participating in extracurricular training and/or activities. If participants answered yes, they were asked to describe the barriers in an open-text field. We categorized the responses into themes. Thirty-nine percent of professionals and 46% of trainees reported facing barriers (Fig. 5). The most commonly cited barriers (by category) were time/workload, supervisor, and awareness/availability/location of activities (Fig. 5). The verbatim responses provided by participants are listed in Supplementary Information 3.

**Fig. 5.**
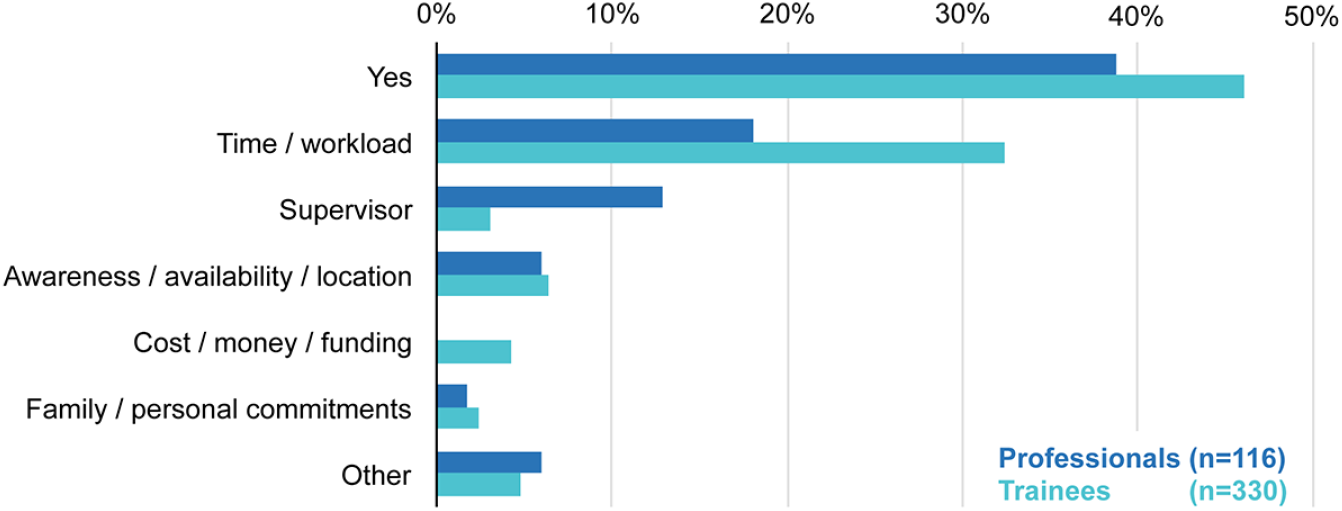
Barriers to participation in extracurricular training and/or activities. **Professionals:** Participants were asked if during their academic training they faced any barriers to participating in extracurricular training and/or activities. **Trainees:** Participants were asked if they face any barriers to participating in extracurricular training and/or activities. **All:** If participants answered yes, they were asked to describe the barriers in an open-text field. Responses were categorized as shown. ‘Other’ includes work environment/culture, unsure of benefits, social anxiety, resources, health issues/disability, qualifications, and other. Refer to Supplementary Information 3 for verbatim responses.

### Value of an advanced degree

Given the investment of time and effort required to complete a PhD and/or postdoctoral fellowship, we wanted to understand how doctoral recipients and former postdoctoral fellows assessed the value of their advanced training. A majority (82%) of professionals (PhD graduates and completed postdoctoral fellows) considered that their PhD helped secure their first position (Fig. 6). Also, most professionals responded that their PhD contributed to their ability to perform at work, and that having a PhD increases their long-term potential for advancement in their chosen field. Eighty-two percent were happy that they pursued a PhD. Among professionals who completed postdoctoral training, 43% thought their postdoctoral training helped start and/or advance their careers, and 51% were happy that they pursued a postdoctoral fellowship (Fig. 6).

**Fig. 6.**
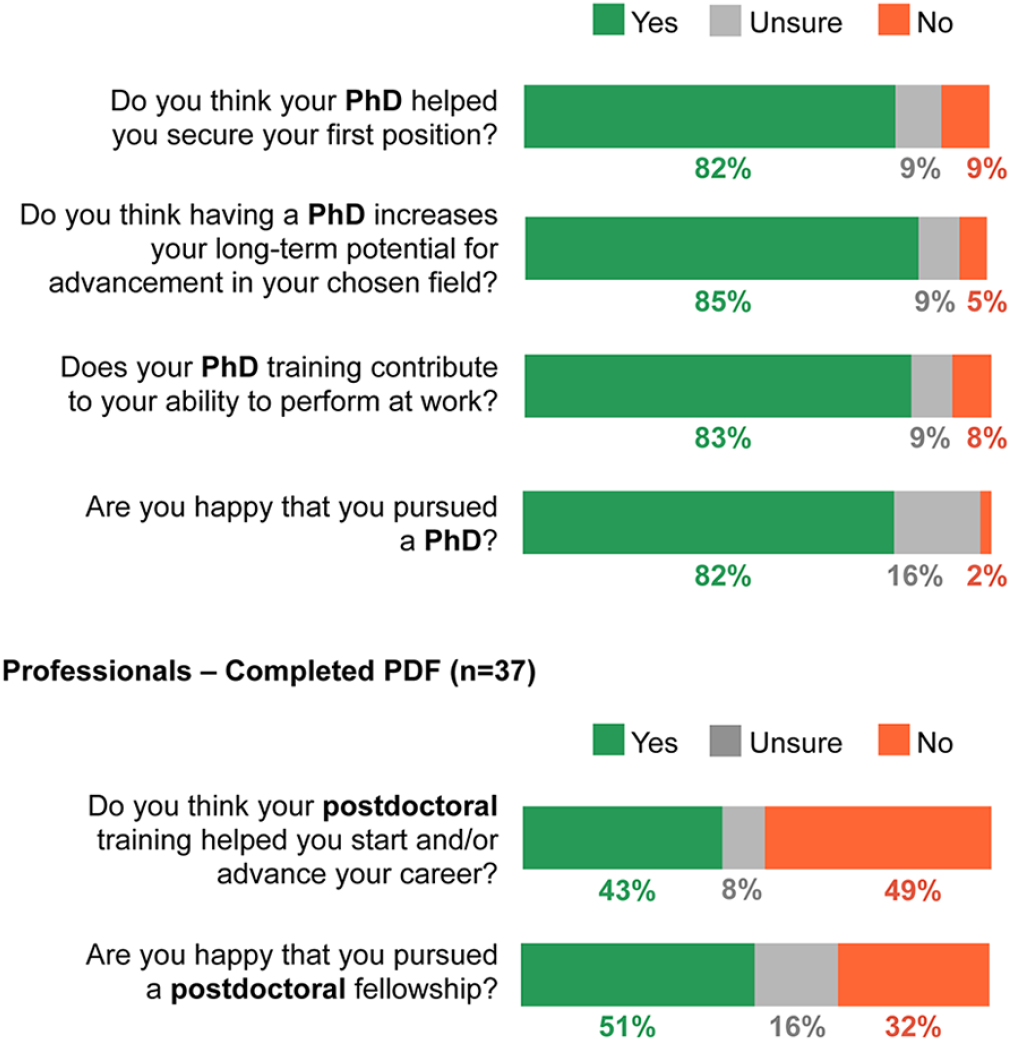
Value and perception of PhD and postdoctoral training. Professionals were asked the questions as shown.

## DISCUSSION

Through this survey we collected the views and perceptions of life science PhD graduates and completed postdoctoral fellows on how career development activities in addition to advanced scientific training have enabled entry into various non-academic careers. We also documented the engagement of life science trainees in career development activities, the barriers encountered by some to participation in such activities, and the rising interest in non-academic careers as trainees approach the completion of their training.

The University of Toronto and the region of Toronto represent a suitable catchment area for this study. The Greater Toronto Area is home to the largest biomedical sciences sector in Canada, including a range of life science research and business opportunities, and academic and non-academic career options (City of Toronto 2018). The University of Toronto has the highest graduate enrolment in Canada (Universities Canada 2017) and the highest ranking in research impact and productivity (Times Higher Education 2018). Given these characteristics, we expect that the survey captured the perceptions and attitudes of talented trainees and graduates from across life science and biomedical departments (Supplementary Information 2).

Our finding of a higher level of interest in non-academic careers among trainees at later stages of their academic programs (Fig. 2b) is consistent with reports about the changing career choices of North American research trainees (Fuhrmann et al. 2011; Sauermann and Roach 2012; Gibbs et al. 2014). A 2010 survey of 39 tier-one U.S. research universities demonstrated significantly decreased preference for academic careers over the course of a PhD program despite active encouragement by thesis advisors for an academic career (Sauermann and Roach 2012). A 2014 survey of 1500 recent American biomedical science PhD graduates demonstrated significantly decreased interest in faculty careers at research universities as well as significantly increased interest in non-research careers at program completion relative to program entry (Gibbs et al. 2014).

The reported drivers of these changes include personal values for work with tangible impact and practical application, and challenging workforce dynamics (e.g., postdoc pay, grant funding, academic job market, etc.) (Gibbs et al. 2013). Despite a shift towards non-academic employment, doctoral and postdoctoral training continues to be dominated by programs that aim to prepare trainees for academic positions (Tilghman et al. 2012).

Professionals in our survey reported high rates of satisfaction with their PhD training, suggesting that advanced training is valuable for a variety of non-academic roles (Fig. 2a, Fig. 6). Relatively fewer professionals who completed postdoctoral fellows rated postdoctoral training favorably (Fig. 6). While a postdoctoral fellowship is often an indispensable step in the academic career path, it may not offer the same value for individuals interested in non-research-based positions outside of academia (Powell 2015; Kahn and Ginther 2017).

While advanced scientific training prepares students for a variety of careers, the traditional, laboratory-based doctoral program may not be sufficient to equip trainees with all of the necessary skills and relatable experiences to compete in today’s non-academic job market. Our findings document trainees’ and professionals’ perception that career development activities beyond the primary laboratory setting play an important and, in some cases, essential role in launching a non-academic career (Fig. 3).

Most survey participants had engaged in at least one career development activity, and participation rates in most activities were comparable between professionals and trainees. These results suggest that many trainees recognize the importance of at least some engagement in career preparation and have taken the initiative to prepare themselves. Activities such as career seminars and networking events had the highest participation rates likely because career exploration is usually an early step undertaken by trainees seeking professional development (Fig. 3). Several activities rated as most impactful, including internship, engagement in not-for-profit organizations or student groups, consulting and entrepreneurship, typically involve experiential learning. Notably, several activities rated by professionals as highly impactful, such as internship, certificate/accreditation, interview training and consulting, had the lowest participation rates among trainees (less than 20%; Fig. 4). There may be many factors underlying this disparity including barriers to participation. Nearly half of all respondents reported facing at least one barrier to participating in extracurricular training and/or activities. These barriers related to both personal factors and the training environment, and included lack of time, lack of awareness, lack of opportunities outside or within graduate programs, and the awareness or perception that some thesis supervisors do not support such activities (Supplementary Information 3). We also acknowledge the temporal disconnect between the pre-graduation experience of professionals and the recent experience of trainees; the perceived value of past activities as rated by professionals may or may not translate to the needs of trainees today.

The activities of perceived high-impact identified in our study reflect other findings in the literature. The importance of experiential learning opportunities such as internships and other work experiences has been highlighted by several authors. For example, a survey of Australian PhD graduates identified paid employment in the final year of doctoral studies as one of the key factors influencing employment after graduation (Jackson and Michelson 2015). Also, a U.K. study found that 80% of doctoral students with work experience reported that it significantly impacted their career decision-making (Vitae 2012). This study also reported the lowest level of work experience during doctoral studies for students in physical, biological and biomedical sciences compared to those in other disciplines. A survey of biomedical postdoctoral fellows in the U.S. demonstrated that while institutions offered increased exposure to career-related information in relation to their graduate studies, the increased knowledge about career options did not translate to increased clarity in their career goals (Gibbs et al. 2015). Based on these findings, the authors suggested that structured career development and exploration opportunities (i.e., experiential learning) are needed in addition to the traditional career development activities such as seminars and panels that provide information on career paths.

While some life science and biomedical departments have begun to incorporate career development training and activities as part of the standard curriculum (e.g., <u>www.nihbest.org</u>; Lee and Reithmeier 2013; Schnoes et al. 2018; University of Toronto Entrepreneurship n.d., University of Calgary 2018), such opportunities do not appear to be common. In contrast, many graduate programs in physical sciences, engineering and computer science actively encourage and accommodate professional experience and entrepreneurial activities (e.g., McMaster University School of Graduate Studies 2017; University of Toronto Entrepreneurship n.d.).

To continue to attract and develop high-potential trainees, faculty and academic administrators may benefit from increasing their awareness of non-academic career paths, encouraging and empowering trainees to explore their interests, and considering how they can support a holistic training curriculum that covers both laboratory research and career development experiences. Other authors have also highlighted the importance of an inclusive, high-quality and satisfying training environment (Duke and Denicolo 2017; Fuhrmann 2016). Various types of career development programming would be relevant to both academic and non-academic employment, and could be designed to allow individuals to tailor their participation to suit their career goals. Also, incoming graduate students and postdoctoral fellows may benefit from asking potential supervisors during the interview process about their support for career development activities. Given the increased interest in non-academic employment closer to completion of training (Fig. 2b), even trainees who are not interested in non-academic employment at the start of their programs could consider probing this topic.

Given that a primary goal of research is to make discoveries and report them, supervisors and trainees may experience tension between research productivity and trainee career preparation. Directly addressing faculty concerns through focus groups, informational meetings, or asking for faculty sign-off on participation in career development training can decrease resistance and strengthen career development programs (Meyers et al. 2016). Profiling faculty members who support both the academic and non-academic career development of their trainees while maintaining research productivity may contribute to the discussion of best-practice. Data on the impact of participating in career development training on time to degree completion and research output are scarce, and further study is warranted.

Our study has several limitations. The survey captured only a subset of the target population, and participants self-selected for meeting the inclusion criteria. Data were collected from eligible trainees and alumni at the University of Toronto and may not be generalizable to all trainees and professionals in life sciences at other institutions. The results include the self-reported attitudes and perceptions of survey participants, which may have been impacted by variations in the content, quality and delivery of extracurricular activities. The study was not designed to compare the career preparation of doctoral degree holders employed outside academia to those employed in academia. Also, we, the authors, have been engaged in developing and delivering career development training and experiences through our volunteer organizations (LSCDS and Science Career Impact Project). Despite these limitations, we believe the findings are worthy of consideration, and we encourage further study on the impact of professional development activities on the employment of doctoral and postdoctoral trainees.

In summary, we found that trainees and professionals employed in non-academic sectors perceived career development activities to play an important role in their path to employment. However, there remains a mismatch between the reported impact of career development activities and participation rates, and many trainees face barriers to pursuing career development activities. Notably, most non-academic professionals value the PhD degree. Our research suggests the need for greater institutional awareness of and support for non-academic career development programs, and increased participation of interested trainees in such programs.

## Supporting information

## Acknowledgements

The authors acknowledge the support of the Graduate Life Sciences Education and Advancement offices at the Faculty of Medicine, University of Toronto, as well as research institutes affiliated with the University of Toronto. This study was funded by the Science Career Impact Project and the Life Sciences Career Development Society.

## Notes

**Potential conflicts of interest:** This study was conducted on a volunteer basis by the authors. The authors have been engaged in developing and delivering career development training and experiences through the Life Sciences Career Development Society and the Science Career Impact Project. No promotion of these programs to the exclusion of other similar programs should be construed. The authors are students and/or alumni of the University of Toronto. MJ and DS are employed in the pharmaceutical industry; their employer was not involved in this work.

