## Supplementary material for "Non-academic employability of life science PhDs: the importance of training beyond the bench"

**Supplementary Information**

**2018**

Sohyoung Her[1,2,*], Mathieu D Jacob[3,*], Sharon Wang[2,*], Songyi Xu[2,4,*], David CF Sealey[3,†,*]

[1] Department of Pharmaceutical Sciences, University of Toronto; [2] Life Sciences Career Development Society, University of Toronto; [3] Science Career Impact Project; [4] Department of Laboratory Medicine and Pathobiology, University of Toronto; Toronto, Ontario, Canada;
* The authors contributed equally to the work.

### 1. Survey

The following survey was developed, and deployed on the internet using SurveyGizmo.

Title: Path to employment for life science PhDs

| ID: 38 |
|---|

We are studying the training and activities that life science trainees at PhD and post-doctoral levels pursue to enhance their employability.

The following cohorts are invited to complete this survey:

- Current life science PhD students and post-doctoral fellows at U of T and affiliated research institutes;
- Individuals who completed life science PhD programs and/or post-doctoral fellowships at U of T and affiliated research institutes after January 1, 2010 AND are employed in a non-academic sector.

The survey is anonymous and will take 10 minutes to complete (on average).

Participants who complete the survey and meet the eligibility criteria described above can enter a random draw for 1 of 3 VISA gift cards valued at $100 each.

If sufficient data are gathered, we may seek to publish aggregate results. Aggregate results may be shared with individual departments and institutes.

Thank you,
Life Science Career Development Society
Science Career Impact Project

| ID: 39 |
|---|

1) What is the highest level of training you have achieved?*
( ) Ph.D. Candidate
( ) Ph.D. Graduate
( ) Current Post-Doctoral Fellow
( ) Completed Post-Doctoral Fellow

| ID: 184 |
|---|

2) What other degrees do you have? * _________________________________________________

Random Draw

Validation: %s format expected
ID: 387

3) If you wish to enter the random draw for the gift card, please provide your e-mail address. Otherwise, proceed to the next question. Your e-mail address will NOT be linked to your responses when the authors analyze the data. Your e-mail address will not be shared with any third party. ________________________________________________

Page entry logic: This page will show when: Question "What is the highest level of training you have achieved?" #1 is one of the following answers ("Ph.D. Candidate","Current Post-Doctoral Fellow")

ID: 407

4) Have you participated in the following types of training? Check all that apply.*
[ ] Resume and/or cover letter training
[ ] Interview training
[ ] Aptitude testing (e.g. personality, work style, strengths)
[ ] Courses outside of field of study (e.g. business, languages, IP)
[ ] Professional skills training (e.g. Project management, negotiation, presentations)
[ ] Certificate/accreditation (e.g. CCRP, CFA, Regulatory)
[ ] Independent study/reading outside of field of study (e.g. trade magazines, blogs, books)
[ ] None of the above

Page entry logic: This page will show when: Question "What is the highest level of training you have achieved?" #1 is one of the following answers ("Ph.D. Candidate","Current Post-Doctoral Fellow")

ID: 399
Piping: Piped From Question 4. (Have you participated in the following types of training? Check all that apply.)

What impact do you think these activities will have on your path to employment?*

| | Highly positive | Positive | Slightly positive | No impact | Unsure |
|---|---|---|---|---|---|

Page entry logic: This page will show when: Question "What is the highest level of training you have achieved?" #1 is one of the following answers
("Ph.D. Candidate","Current Post-Doctoral Fellow")

ID: 409

5) Have you participated in any of the following activities? Check all that apply.*
[ ] Career seminars
[ ] Networking events
[ ] Company information sessions/site visits
[ ] Conferences outside field of study
[ ] Informational interviews
[ ] Mentorship/career coaching
[ ] Involvement in student or non-profit organization
[ ] Case competitions
[ ] Entrepreneurship
[ ] Consulting (paid or volunteer)
[ ] Internship (paid or volunteer)
[ ] Research/publication outside of field of study
[ ] None of the above

Page entry logic: This page will show when: Question "What is the highest level of training you have achieved?" #1 is one of the following answers
("Ph.D. Candidate","Current Post-Doctoral Fellow")

ID: 410
Piping: Piped From Question 5. (Have you participated in any of the following activities? Check all that apply.)

What impact do you think these activities will have on your path to employment?*

| | Highly positive | Positive | Slightly positive | No impact | Unsure |
|---|---|---|---|---|---|

Page entry logic: This page will show when: Question "What is the highest level of training you have achieved?" #1 is one of the following answers ("Ph.D. Candidate","Current Post-Doctoral Fellow")

ID: 420

6) Have you participated in any other extra-curricular training or activities? If yes, please identify the activities and the impact you think they will have on your path to employment.*

________________________________________________

ID: 419

7) Do you face any barriers to participating in extra-curricular training and/or activities? If yes, please describe.*

( ) Yes: _____________________________________________________*
( ) No

ID: 174

8) Do you plan to engage in the following extra-curricular training or activities in the next 12 months? Please select all that apply.*

[ ] Resume and/or cover letter training
[ ] Interview training
[ ] Aptitude testing (e.g. personality, work style, strengths)
[ ] Courses outside of field of study (e.g. business, languages, IP)
[ ] Professional skills training (e.g. Project management, negotiation, presentations)
[ ] Certificate/accreditation (e.g. CCRP, CFA, Regulatory)
[ ] Independent study/reading outside of field of study (e.g. trade magazines, blogs, books)
[ ] Career seminars
[ ] Networking events
[ ] Company information sessions/site visits
[ ] Conferences outside field of study
[ ] Informational interviews
[ ] Mentorship/career coaching
[ ] Involvement in student or non-profit organization
[ ] Case competitions
[ ] Entrepreneurship
[ ] Consulting (paid or volunteer)
[ ] Internship (paid or volunteer)
[ ] Research/publication outside of field of study
[ ] Other: _____________________________________________________*
[ ] None of the above

Page entry logic: This page will show when: Question "What is the highest level of training you have achieved?" #1 is one of the following answers
("Ph.D. Graduate","Completed Post-Doctoral Fellow")

ID: 413

9) During your academic training, did you participate in the following types of training? Check all that apply.*
[ ] Resume and/or cover letter training
[ ] Interview training
[ ] Aptitude testing (e.g. personality, work style, strengths)
[ ] Courses outside of field of study (e.g. business, languages, IP)
[ ] Professional skills training (e.g. Project management, negotiation, presentations)
[ ] Certificate/accreditation (e.g. CCRP, CFA, Regulatory)
[ ] Independent study/reading outside of field of study (e.g. trade magazines, blogs, books)
[ ] None of the above

Page entry logic: This page will show when: Question "What is the highest level of training you have achieved?" #1 is one of the following answers
("Ph.D. Graduate","Completed Post-Doctoral Fellow")

ID: 414
Piping: Piped From Question 9. (During your academic training, did you participate in the following types of training? Check all that apply.)

What impact did these activities have on your path to employment?*

| | Highly positive - no job without it | Positive - it was an asset | Slightly positive - nice to have | No impact | Unsure |
|---|---|---|---|---|---|

Page entry logic: This page will show when: Question "What is the highest level of training you have achieved?" #1 is one of the following answers
("Ph.D. Graduate","Completed Post-Doctoral Fellow")

ID: 415

10) During your academic training, did you participate in any of the following activities? Check all that apply.*
[ ] Career seminars
[ ] Networking events
[ ] Company information sessions/site visits
[ ] Conferences outside field of study
[ ] Informational interviews
[ ] Mentorship/career coaching
[ ] Involvement in student or non-profit organization
[ ] Case competitions
[ ] Entrepreneurship
[ ] Consulting (paid or volunteer)
[ ] Internship (paid or volunteer)
[ ] Research/publication outside of field of study
[ ] None of the above

Page entry logic: This page will show when: Question "What is the highest level of training you have achieved?" #1 is one of the following answers
("Ph.D. Graduate","Completed Post-Doctoral Fellow")

ID: 416
Piping: Piped From Question 10. (During your academic training, did you participate in any of the following activities? Check all that apply.)

What impact did these activities have on your path to employment?*

| | Highly positive - no job without it | Positive - it was an asset | Slightly positive - nice to have | No impact | Unsure |
|---|---|---|---|---|---|

Page entry logic: This page will show when: Question "What is the highest level of training you have achieved?" #1 is one of the following answers
("Ph.D. Graduate","Completed Post-Doctoral Fellow")

ID: 343

11) During your academic training, did you participate in any other extra-curricular training or activities that facilitated your path to employment? If yes, please identify the activities and their impact on your path to employment.* ____________________________

ID: 418

12) During your academic training, did you face any barriers to participating in extra-curricular training and/or activities? If yes, please describe.*
( ) Yes: _________________________________________________*
( ) No

ID: 346

13) Do you think your PhD helped you secure your first position?*
( ) Yes
( ) No
( ) Unsure

ID: 417

14) After you finished your academic training, how long did it take you to secure a position that you were professionally satisfied with?*
( )
( ) 1 year
( ) 2 years
( ) 3 years
( ) 4 years
( ) 5 years
( ) >5 years
( ) Still working on it

ID: 349
15) Does your PhD training contribute to your ability to perform at work?*
( ) Yes
( ) No
( ) Unsure

ID: 350

16) Do you think having a PhD increases your long-term potential for advancement in your chosen field?*
( ) Yes
( ) No
( ) Unsure

ID: 352

17) Are you happy that you pursued a PhD?*
( ) Yes
( ) No
( ) Unsure

Page entry logic: This page will show when: Question "What is the highest level of training you have achieved?" #1 is one of the following answers
("Completed Post-Doctoral Fellow")

ID: 383

18) Do you think your postdoctoral training helped you start and/or advance your career?*
( ) Yes
( ) No
( ) Unsure

ID: 385

19) Are you happy that you pursued a postdoctoral fellowship?*
( ) Yes
( ) No
( ) Unsure

Page entry logic: This page will show when: Question "What is the highest level of training you have achieved?" #1 is one of the following answers ("Ph.D. Candidate","Current Post-Doctoral Fellow")

ID: 180

20) What sector do you wish to work in?*
( ) Non-academic (e.g., Industry, government, non-profit)
( ) Academic (e.g., university, research institute)
( ) Other: _________________________________________________*

ID: 181

21) When do you plan to finish your training?*
( ) 2017
( ) 2018
( ) 2019
( ) 2020
( ) 2021 or beyond

ID: 394

22) Which academic department are you affiliated with?*
( ) Biochemistry
( ) Cell & Systems Biology
( ) Immunology
( ) Institute of Biomaterials and Biomedical Engineering
( ) Institute of Medical Sciences
( ) Laboratory Medicine and Pathobiology
( ) Medical Biophysics
( ) Molecular Genetics
( ) Nutritional Sciences
( ) Pharmaceutical Sciences
( ) Pharmacology and Toxicology
( ) Physiology
( ) Other: _________________________________________________*

ID: 395

23) Which institution are you affiliated with?*
( ) Bloorview Research Institute
( ) CAMH
( ) Donnelly Centre for Cellular and Biomolecular Research
( ) Hospital for Sick Children
( ) Keenan Research Centre for Biomedical Science, St. Michael's Hospital
( ) Kunin-Lunenfeld Applied & Evaluative Research Unit, Baycrest
( ) Li Ka Shing Knowledge Institute, St. Michael's Hospital
( ) Lunenfeld-Tanenbaum Research Institute, Mount Sinai Hospital
( ) McLaughlin Centre for Molecular Medicine
( ) Rotman Research Institute
( ) Sunnybrook Research Institute
( ) Tanz Centre for Research in Neurodegenerative Diseases
( ) University Health Network (TGRI, PMH, Western)
( ) University of Toronto, Mississauga
( ) University of Toronto, Scarborough
( ) Women's College Research Institute
( ) Other: _________________________________________________*

ID: 194

24) What is the focus area of your research?*
( ) Basic Research
( ) Applied Research
( ) Clinical Research
( ) Other: _________________________________________________*

Page entry logic: This page will show when: Question "What is the highest level of training you have achieved?" #1 is one of the following answers
("Ph.D. Graduate","Completed Post-Doctoral Fellow")

ID: 396

25) In which department did you complete your academic training?*
( ) Biochemistry
( ) Cell & Systems Biology
( ) Immunology
( ) Institute of Biomaterials and Biomedical Engineering
( ) Institute of Medical Sciences
( ) Laboratory Medicine and Pathobiology
( ) Medical Biophysics
( ) Molecular Genetics
( ) Nutritional Sciences
( ) Pharmaceutical Sciences
( ) Pharmacology and Toxicology
( ) Physiology
( ) Other: _________________________________________________*

ID: 397

26) What was your institutional affiliation?*
( ) Bloorview Research Institute
( ) CAMH
( ) Donnelly Centre for Cellular and Biomolecular Research
( ) Hospital for Sick Children
( ) Keenan Research Centre for Biomedical Science, St. Michael's Hospital
( ) Kunin-Lunenfeld Applied & Evaluative Research Unit, Baycrest
( ) Li Ka Shing Knowledge Institute, St. Michael’s Hospital
( ) Lunenfeld-Tanenbaum Research Institute, Mount Sinai Hospital
( ) McLaughlin Centre for Molecular Medicine
( ) Rotman Research Institute
( ) Sunnybrook Research Institute
( ) Tanz Centre for Research in Neurodegenerative Diseases
( ) University Health Network (TGRI, PMH, Western)
( ) University of Toronto, Mississauga
( ) University of Toronto, Scarborough
( ) Women's College Research Institute
( ) Other: _________________________________________________*

ID: 287

27) What was the focus area of your research?*
( ) Basic Research
( ) Applied Research
( ) Clinical Research
( ) Other: _________________________________________________*

Page entry logic: This page will show when: Question "What is the highest level of training you have achieved?" #1 is one of the following answers
("Ph.D. Graduate")

ID: 282

28) What was the duration of your PhD?*
( ) 3 years
( ) 4 years
( ) 5 years
( ) 6 years
( ) 7 years
( ) 8 years
( ) 9+ years

ID: 283

29) How many years has it been since you completed your PhD?*
( ) 1 year
( ) 2 years
( ) 3 years
( ) 4 years
( ) 5 years
( ) 6 years
( ) 6+ years

Page entry logic: This page will show when: Question "What is the highest level of training you have achieved?" #1 is one of the following answers
("Completed Post-Doctoral Fellow")

ID: 220

30) What was the duration of your post-doctoral training period?*
( ) 1 year
( ) 2 years
( ) 3 years
( ) 4 years
( ) 5 years
( ) 6 years
( ) 7 years
( ) 8 years
( ) 9+ years

ID: 221

31) How many years has it been since you completed your postdoctoral training?*
( ) 1 year
( ) 2 years
( ) 3 years
( ) 4 years
( ) 5 years
( ) 6 years
( ) 6+ years

Page entry logic: This page will show when: Question "What is the highest level of training you have achieved?" #1 is one of the following answers
("Ph.D. Graduate","Completed Post-Doctoral Fellow")

ID: 266

32) What sector do you currently work in?*
( ) Non-academic (e.g., Industry, government, non-profit)
( ) Academic (e.g., university, research institute)
( ) Other: _________________________________________________*

ID: 268

33) What functional areas or departments (e.g., R&D, policy, business development, medical affairs) have you worked in?* _________________________________________________

Page entry logic: All respondents

Final questions

ID: 185

34) Is there anything else you would like the study authors to know about your training and/or career path? _________________________________________________

Validation: %s format expected
ID: 398

35) If you would you be willing to answer additional questions about your responses, please provide your e-mail address. Your e-mail address will be linked to your responses. Your e-mail address will not be shared with any third party. ____________________________________
Thank You!

ID: 1

Thank you for completing the survey.

### 2. University of Toronto department affiliation

PhD candidate and post-doctoral fellow survey participants were asked Question 22: "Which academic department are you affiliated with?" PhD graduate and completed post-doctoral fellow survey participants were asked Question 25: "In which department did you complete your academic training?" Responses are tabulated below.

| | Trainees | | Professionals | | |
|---|---|---|---|---|---|
| | PhD Candidates n=244 | Postdoctoral Fellows n=86 | Completed PhD n=79 | Completed Postdoctoral Fellowship n=37 | Total N=446 |
| Molecular Genetics | 34 (13.9%) | 12 (14.0%) | 16 (20.3%) | 11 (29.7%) | 73 (16.4%) |
| Medical Biophysics | 22 (9.0%) | 11 (12.8%) | 14 (17.7%) | 6 (16.2%) | 53 (11.9%) |
| Institute of Medical Sciences | 25 (10.2%) | 5 (5.8%) | 8 (10.1%) | 3 (8.1%) | 41 (9.2%) |
| Laboratory Medicine & Pathobiology | 15 (6.1%) | 9 (10.5%) | 8 (10.1%) | 2 (5.4%) | 34 (7.6%) |
| Cell & Systems Biology | 22 (9.0%) | 9 (10.5%) | 0 (0.0%) | 1 (2.7%) | 32 (7.2%) |
| Immunology | 15 (6.1%) | 6 (7.0%) | 6 (7.6%) | 2 (5.4%) | 29 (6.5%) |
| Biochemistry | 12 (4.9%) | 4 (4.7%) | 6 (7.6%) | 5 (13.5%) | 27 (6.1%) |
| Physiology | 15 (6.1%) | 4 (4.7%) | 2 (2.5%) | 2 (5.4%) | 23 (5.2%) |
| Inst. of Biomaterials & Biomed. Eng. | 12 (4.9%) | 3 (3.5%) | 3 (3.8%) | 2 (5.4%) | 20 (4.5%) |
| Pharmaceutical Sciences | 11 (4.5%) | 2 (2.3%) | 6 (7.6%) | 0 (0.0%) | 19 (4.3%) |
| Pharmacology & Toxicology | 11 (4.5%) | 2 (2.3%) | 4 (5.1%) | 1 (2.7%) | 18 (4.0%) |
| Rehabilitation Sciences Institute | 11 (4.5%) | 1 (1.2%) | 0 (0.0%) | 0 (0.0%) | 12 (2.7%) |
| Ecology & Evolutionary Biology | 10 (4.1%) | 1 (1.2%) | 0 (0.0%) | 0 (0.0%) | 11 (2.5%) |
| Nutritional Sciences | 8 (3.3%) | 2 (2.3%) | 0 (0.0%) | 1 (2.7%) | 11 (2.5%) |
| Chemistry | 8 (3.3%) | 0 (0.0%) | 2 (2.5%) | 0 (0.0%) | 10 (2.2%) |
| Psychology | 6 (2.5%) | 3 (3.5%) | 1 (1.3%) | 0 (0.0%) | 10 (2.2%) |
| Public Health | 2 (0.8%) | 2 (2.3%) | 0 (0.0%) | 0 (0.0%) | 4 (0.9%) |
| Chemical Engineering | 2 (0.8%) | 0 (0.0%) | 0 (0.0%) | 0 (0.0%) | 2 (0.4%) |
| Neuroscience | 1 (0.4%) | 1 (1.2%) | 0 (0.0%) | 0 (0.0%) | 2 (0.4%) |
| Inst. of Health Policy, Mgt. & Eval. | 0 (0.0%) | 2 (2.3%) | 0 (0.0%) | 0 (0.0%) | 2 (0.4%) |
| Psychiatry | 0 (0.0%) | 2 (2.3%) | 0 (0.0%) | 0 (0.0%) | 2 (0.4%) |
| Cognitive Neuropsychology | 1 (0.4%) | 0 (0.0%) | 0 (0.0%) | 0 (0.0%) | 1 (0.2%) |
| Kinesiology | 1 (0.4%) | 0 (0.0%) | 0 (0.0%) | 0 (0.0%) | 1 (0.2%) |
| Genetics & Development | 0 (0.0%) | 1 (1.2%) | 0 (0.0%) | 0 (0.0%) | 1 (0.2%) |
| Inst. for Mental Health Policy Research | 0 (0.0%) | 1 (1.2%) | 0 (0.0%) | 0 (0.0%) | 1 (0.2%) |
| Matrix Dynamics | 0 (0.0%) | 1 (1.2%) | 0 (0.0%) | 0 (0.0%) | 1 (0.2%) |
| Nursing | 0 (0.0%) | 1 (1.2%) | 0 (0.0%) | 0 (0.0%) | 1 (0.2%) |
| Experimental Therapeutics | 0 (0.0%) | 1 (1.2%) | 0 (0.0%) | 0 (0.0%) | 1 (0.2%) |
| Epidemiology | 0 (0.0%) | 0 (0.0%) | 1 (1.3%) | 0 (0.0%) | 1 (0.2%) |
| Faculty of Dentistry | 0 (0.0%) | 0 (0.0%) | 1 (1.3%) | 0 (0.0%) | 1 (0.2%) |
| Microbiology | 0 (0.0%) | 0 (0.0%) | 1 (1.3%) | 0 (0.0%) | 1 (0.2%) |
| Oncology | 0 (0.0%) | 0 (0.0%) | 0 (0.0%) | 1 (2.7%) | 1 (0.2%) |

### 3. Barriers to participating in extra-curricular training and/or activities

PhD candidate and post-doctoral fellow survey participants were asked Question 7: “Do you face any barriers to participating in extra-curricular training and/or activities?” PhD graduate and completed post-doctoral fellow survey participants were asked Question 12: “During your academic training, did you face any barriers to participating in extra-curricular training and/or activities?” If participants answered yes, they were asked to describe the barriers in an open-text field.

Responses were categorized as follows: time/workload; supervisor; awareness/availability/ location; cost/money/funding; family/personal commitments; other. If applicable, responses were counted in more than one category. Category frequencies are shown in Figure 5.

The verbatim responses entered by participants into the open-text field are provided in the listings below in alphabetical order per category (next page). For responses counted in more than one category, the responses appear in each of the applicable categories. Responses provided by more than one participant are indicated by “(multiple).”

Barrier category: **Time/workload**
Respondent type: **Trainees – PhD candidates**

Always very busy in the lab

Amount of time available outside of research and teaching

Between school and work it is difficult to find time.

Conflict of time

Difficult to allocate time away from lab, many events run at UofT are during regular work hours, would be good to have evening events as well

Financial - I work part-time in addition to full-time graduate studies to support my household

Financial, time

Financial, time limitations

Guilt that I'm taking time away from my research

Inadequate time

It is hard to find time

Lab commitment (9-5)

Lab work (multiple)

Lab Work Needs to be Done - not a complaint, just a fact

Lab work schedule

Lab workload

Lack of awareness, lack of time

Lack of time (multiple)

Lack of time, difficult to find

Lack of time, health concerns

Limited time (multiple)

Location, time, so mostly conflict with lab/academic research time

My own study, lack of time to focus on extracurricular

Need to spend more time in lab

No time

No Time!

No time/money

No time outside of lab

Not enough time

Not much time left in the day!

PhD is stressful, demanding; time availability depends a lot on project, PI...

Requirements in the psych program are greater than most - making it difficult to find time to do extra-curriculars

Scheduling conflicts with graduate study commitments

Sometimes there is a lack of time

Takes time away from school

Taking time away from research/goal of finishing

Time (multiple)

Time! (multiple)

Time and anxiety in meeting new people

Time and money (multiple)

Time and money!

Time. And varying amounts of support from supervisor.

Time availability

Time away from lab work

Time away from the lab

Time commitment (multiple)

Time commitment, cost

Time constraint

Time constraints

Time constraints. I have found that as you become more senior in your lab you are required to write more and more grants, lit reviews, editorials, review manuscripts, etc., so you already have less time for your own work, let alone extra-curriculars.

Time constraints with graduate program

Time consuming (multiple)

Time in the lab

Time is often limited, and they are often expensive.

Time! I think the biggest problem is balancing the time for course work/TA-ing/research-lab work, leaving little time for extracurricular activities unless it becomes a necessity.

Time (lack thereof)

Time management (multiple)

Time more than anything else. It is difficult to juggle extra-curricular training with a thesis requirements and family.

Time - most of my time has to be spent in lab

Time (time away from graduate work/thesis)

Time, unable to walk away from the lab

Time, undergraduate GPA

Tricky to take time off of lab work for competitions etc

Barrier category: **Time/workload**

| **Trainees** | | **Professionals** | |
|---|---|---|---|
| **PhD candidates** | **Postdoctoral fellows** | **PhD graduates** | **Completed postdoctoral fellows** |
| (see previous page) | Amount of time available outside of research and teaching | At times difficult to balance lab work and extracurriculars | No time |
| | Amount of work to be done in my own research projects | Clinical studies demand | Shortage of Time |
| | As a full time postdoc the expectation is to even engage your personal time on this activity, which does not leave much time to explore alternatives to academy | Hard to find time | Time commitment outside research is difficult |
| | Between school and work it is difficult to find time. | Lack of time | Too busy, not aware, unsure of benefits, narrow focus |
| | Conflict of time | Lack of time and resources | |
| | It's difficult to find opportunities and the time. | May be viewed as taking too much time away from academic/research work | |
| | Lack of time | No time | |
| | Little time, and less reward. Not recognized enough. | Not enough time and mentorship from academics | |
| | Managing time and prioritizing are always challenging | Not enough time for these activities | |
| | No time | Time (multiple) | |
| | No time/money | Time away from lab | |
| | Not enough time | Time constraints | |
| | Overwhelming amount of lab work. | Time constraints due to raising a family | |
| | Time (multiple) | Too little time left after research work | |
| | Time, due to research | | |
| | Time off of lab work for competitions etc | | |

Barrier category: **Supervisor**

### Trainees

#### PhD candidates

- My supervisor looks down upon it.
- PhD is stressful, demanding; time availability depends a lot on project, PI...
- PI only values research. Everything professional development activity undertaken must be hidden from PI.
- Sometimes supervisors can be a barrier to participating in activities
- Supervisor discourages activities outside research
- Supervisors discourages extra-curricular activities
- Time. And varying amounts of support from supervisor.

#### Postdoctoral fellows

- Although my doctoral supervisor never stopped me, she didn't particularly like it
- My current boss
- Sometime money is an issue. Also, overwhelming demands from professors who think extra-curricular activities are distractions

### Professionals

#### PhD graduates

- Had to do it in secret - academic supervisor would not have been supportive
- Had to schedule around lab work so PI would not know. (multiple)
- It was not encouraged openly by PI
- My supervisor was not very supportive of my extracurricular activities and never encouraged me to do so
- PhD supervisor did not like any work or activities outside of the lab, had to actively hide the courses that I was taking, and the other extracurriculars
- PhD supervisor was opposed to me spending time outside of my thesis if not required.
- PI's expected me to be in the lab, so it was very difficult to participate in career development events especially during office hours.
- Supervisor not supporting time away from bench
- Time off from PI was limit

#### Completed postdoctoral fellows

- I was not encouraged to do so, so I hid my involvement from my supervisors and most of my lab mates
- My postdoctoral supervisor strongly discouraged working outside the lab (as a course instructor)
- Postdoc mentor discouraged extra-curricular training and/or activities
- Supervisor did not approve of training during regular work hours

Barrier category: **Awareness/availability/location**

| **Trainees** | | **Professionals** | |
|---|---|---|---|
| **PhD candidates** | **Postdoctoral fellows** | **PhD graduates** | **Completed postdoctoral fellows** |
| Commute (multiple) | Availability of such activities | Distance between lab and downtown core, didn't seem encouraged | Lack of awareness |
| Currently not in Toronto | Family and commuting, most events are in the evening and I am unable to attend. | I was involved in many extra-curricular activities, but it was a challenge to become aware of the opportunities available. I had to go out of my way to find them. In particular TAing was something so classmates wanted to get involved in but the opportunities were scarce in my department (Molecular Genetics). | None offered |
| Difficult to allocate time away from lab, many events run at UofT are during regular work hours, would be good to have evening events as well | Hard to find ones relevant to my field | Scheduling limitations | Too busy, not aware, unsure of benefits, narrow focus |
| Finding the opportunities | Having graduated | | |
| Location, time, so mostly conflict with lab/academic research time | It's difficult to find opportunities and the time. | | |
| Lack of advertising for these opportunities | Knowing which are most impactful and therefore worth the time investment | | |
| Lack of awareness, lack of time | Lack of opportunities offered | | |
| Lack of communication | Limited opportunity in Canada (most PhD grads head to the US for exposure) | | |
| Lack of knowledge of most relevant extra-curricular activities | Postdocs are sometimes excluded from such trainings | | |
| Lack of time/difficult to find | | | |
| Poor timing of activities in general (if they are within 9-5) | | | |
| There are many events, but it is hard to determine which ones will be most useful. Some events are circulated very late as well (ie. day of), which makes it difficult to attend Working remotely / 'field research' is isolating from on-campus resources | | | |
| There seem to be fewer opportunities for involvement being a student at UTM | | | |

Barrier category: **Cost/money/funding**

| Trainees | | Professionals | |
|---|---|---|---|
| **PhD candidates** | **Postdoctoral fellows** | **PhD graduates** | **Completed postdoctoral fellows** |
| Financial - I work part-time in addition to full-time graduate studies to support my household | Monetary - sometimes | | |
| Financial, time | Money (multiple) | | |
| Financial / time limitations | Money! | | |
| Funding would be helpful. | No time/money | | |
| No time/money | Sometime money is an issue. Also, overwhelming demands from professors who think extra-curricular activities are distractions | | |
| Time and money (multiple) | | | |
| Time and money! | | | |
| Time commitment, cost | | | |
| Time is often limited, and they are often expensive. | | | |

Barrier category: **Family/personal commitments**

| **Trainees** | | **Professionals** | |
|---|---|---|---|
| **PhD candidates** | **Postdoctoral fellows** | **PhD graduates** | **Completed postdoctoral fellows** |
| Dependent child | | Had a child during PhD | |
| Family and commuting, most events are in the evening and I am unable to attend. | | Time constraints due to raising a family | |
| Family commitments (parenting) | | | |
| Family responsibilities | | | |
| Financial - I work part-time in addition to full-time graduate studies to support my household | | | |
| I have an infant who requires my care outside regular working hours. | | | |
| Often after standard working hours and I have other commitments to tend to | | | |
| Personal life commitments | | | |
| Time more than anything else. It is difficult to juggle extra-curricular training with a thesis requirements and family. | | | |

Barrier category: **Other** *(category)*

| **Trainees** | | **Professionals** | |
|---|---|---|---|
| **PhD candidates** | **Postdoctoral fellows** | **PhD graduates** | **Completed postdoctoral fellows** |
| Disability related issues *(health issue)* | Little time, and less reward. Not recognized enough. *(work environment/culture)* | Completing my academic training *(other)* | I was not encouraged to do so, so I hid my involvement from my supervisors and most of my lab mates *(work environment/culture)* |
| Experienced severe episode of clinical depression for 1.5 years. *(health issue)* | Not enough background to secure spot for miniMBA program *(credentials)* | Distance between lab and downtown core, didn't seem encouraged *(work environment/culture)* | Too busy, not aware, unsure of benefits, narrow focus *(unsure of benefits)* |
| I feel I'm too old in some of these. *(social issue)* | | Lack of time and resources *(resources)* | Pressure to be in the lab at all hours *(work environment/ culture)* |
| Lack of time, health concerns *(health issue)* | | May be viewed as taking too much time away from academic/ research work *(work environment/culture)* | |
| Seems impossible to find relevant mentors via mentorship programs *(resources)* | | Not enough time and mentorship from academics *(work environment/culture)* | |
| Social *(social issue)* | | Other (multiple) | |
| Social anxiety *(social issue)* | | | |
| Time and anxiety in meeting new people *(social issue)* | | | |
| Time, undergraduate GPA *(credentials)* | | | |
| Other | | | |
